# Integrated Immunopeptidomics and Proteomics Study Reveals Imbalanced Innate and Adaptive Immune Responses to SARS-Cov-2 Infection

**DOI:** 10.1101/2022.08.23.504798

**Authors:** Rui Chen, Kelly M. Fulton, Anh Tran, Diana Duque, Kevin Kovalchik, Etienne Caron, Susan M. Twine, Jianjun Li

**Affiliations:** Human Health Therapeutics Research Centre, National Research Council Canada, 100 Sussex Drive, Ottawa, Ontario, Canada, K1A 0R6; CHU Sainte-Justine Research Center, Montreal, QC H3T 1C5, Canada; Department of Pathology and Cellular Biology, Faculty of Medicine, Université de Montréal, QC H3T 1J4, Canada

## Abstract

We present an integrated immunopeptidomics and proteomics study of SARS-Cov-2 infection to comprehensively decipher the changes in host cells in response to viral infection. Our results indicated that innate immune response in Calu-3 cells was initiated by TLR3, followed by activation of interferon signaling pathway. Host cells also present viral antigens to the cell surface through both Class I and Class II MHC system for recognition by adaptive immune system. SARS-Cov-2 infection led to the disruption of antigen presentation as demonstrated by higher level of HLA proteins from the flow-through of MHC immunoprecipitation. Glycosylation analysis of HLA proteins from the elution and flow-through of immunoprecipitation revealed that the synthesis and degradation of HLA protein was affected by SARS-Cov-2 infection. This study provided many useful information to study the host response to SARS-Cov-2 infection and would be helpful for the development of therapeutics and vaccine for Covid-19 and future pandemic.

## Introduction

The novel severe acute respiratory syndrome coronavirus 2 (SARS-CoV-2) is the causative agent of COVID-19. This was declared a pandemic in March 2020. Since then, the world has witnessed the mobilisation of the global scientific community toward the fastest vaccine development in history. Currently there are 14 vaccines approved by WHO for emergent used globally, with 341 in various stages of development. After two years into the pandemic, there is still a need to understand how the virus mobilises the host cellular pathways to allow rapid infection and replication, and the key viral epitopes that stimulate the host immune system to generate protective immunity.

SARS-CoV-2 is a beta coronavirus, a member of the Coronaviridae family of viruses, include four sub-groups, denoted alpha, beta, gamma and delta. Together, the family infect a wide range of mammal and bird species, with mutations allowing the species barrier to be crossed. SARS-CoV-2 has a 30 kb positive-sense RNA genome, with 14 open reading frames (ORFs), encoding a total of 29 proteins. These include structural proteins (Spike, Envelope, Membrane, Nucleocapsid), 16 non-structural proteins (Nsp 1-16) and 9 accessory proteins (Orf 3a,b, Orf6, Orf7a, b, Orf8, Orf 9b, c and Orf 10)^1^. Building on knowledge of related coronaviruses, recent studies have illuminated the lifecycle of SARS-CoV-2 ^2^. This virus enters cells via binding of the spike structural protein to human angiotensin-converting enzyme 2 (ACE2) on the surface of host cells. Intracellularly, the viral genome is transcribed by the host ribosomes. The viral genome is replicated by the viral RNA-dependent RNA polymerase, on membrane structures derived from the endoplasmic reticulum (ER). Assembly of mature virions occurs in the ER-golgi intermediate compartment, budding from the ER-Golgi membranes before release outside the cells via an exocytosis like process^3^. Despite the rapid progress made by the scientific community, the challenge of the emergence of variants of concern (VoC), continues to challenge vaccine and drug developers. Although the current approved vaccines showed excellent efficacy toward the original SARS-CoV-2 virus, vaccine breakthrough infections have been observed with several VoCs. This continues to highlight the need to build on our current knowledge of the host response to the virus, in particular the signalling pathways modified to allow viral infection, and the viral epitopes presented to the immune system.

The genome sequence was widely disseminated early on ^4^ allowing the rapid design of first generation vaccines. Infection with the novel coronavirus elicits both humoral and cell mediated immunity. Although vaccines offer protection against the original virus and some protection in terms of reduced severity of infection and transmission of VoC, there is still a need for booster vaccines and universal vaccines that offer protection from variants of concern. Despite the approval of vaccines, variants of concern continue to emerge. There are also concerns about resistance to antibody defences. There is a need to understand immune responses, such as cytotoxic T-cell could boost immunity ^5^. Viruses infect host cells and their proteins are processed and presented on the surface of human cells by HLA-I. Cytotoxic T-cells recognised non-self antigens and initiate an immune response that clears the infected cells. Improving knowledge of the immunpeptidome will shed light on those peptides that activate cytotoxic T-cells. This will require a knowledge of the viral peptide epitopes presented to the host immune system. Advanced in silico methods are used to predict immune epitopes, but it is not known whether these epitopes confer protective immunity or stimulate a t-cell response. In silico methods are a critical tool, but cannot yet take into account the ways viruses can manipulate host cells processes, impacting antigen presentation. For example, viruses have been shown to downregulate the proteasome, host cell protein translation and HLA-I expression^6^.

Prediction modelling does not take into account kinetics of viral suppression on HLA-I expression, which may mean that there is a larger repertoire of viral epitopes early on during infection of host cells, ^7^. Studies typically use overlapping peptide tiling and/or in silico prediction approaches to forecast HLA-I binding peptides ^8-13^. Whilst in silico prediction provides a richness of information regarding immune epitopes, the information needs to be confirmed experimentally and the identified peptides tested for T-cell activation. For this, mass spectrometric sequencing of peptides is required. Mass spectrometry can be used to identify viral peptides bound to MHC complexes - it is a direct and target method to discover endogenously presented peptides. The approach has been used in other viral infections, revealing new antigens including those peptide sequences that activated T-cell responses against West Nile virus^14^, HIV^15,16^ and measles^17^. It was reported that epitopes of SARS-CoV-2 derived not only from structural but also non-structural genes in regions highly conserved among SARS-CoV-2 strains including recently recognized variants^18^. This included epitopes from membrane glycoprotein (MGP) and non-structure protein-13 (NSP13). A recent study investigated the HLA-I immunopeptidome of two SARS-CoV-2 infected human cells lines, identifying HLA-I bound peptides derived from canonical and non-canonical ORFs^19^. Non-canonical ORFs are not reflected in current vaccines. Other studies have combined HLA peptide sequencing with host cell protein abundance proteomics, which showed that those viral proteins expressed early in infection were found more frequently in HLA presentation ^20^.

In addition to immunopeptidomics, proteomics analysis studies the changes of host proteome after infection can provide more insights into the molecular basics of SARS-Cov-2 infection. Several studies used different cell lines and techniques to study the change in either global proteome or PTM specific subproteome to identify the pathways involved in response to viral infection and develop therapeutics against Covid-19^21-25^. However, immunopeptidomics were not included in those studies while the host proteome change was not studied in depth from several immunopeptidomics analysis^19,26^,27. As shown in recent studies, SARS-Cov-2 infection will hijack the antigen presentation pathways and prevent infected cells from presenting viral antigens ^28,29^, which might cause the impaired T cell response in Covid-19 patients ^30^. Thus, it is more important to study how the infection impacts the antigen presentation, which is vital to the adaptive immune system. To fill this gap, we developed an integrated immunopeptidomics and proteomics approach that uses MHC immune-precipitation, combined with mass spectrometry to survey the SARS-Cov-2 immunopeptidome and changes in the host proteome. In additional to the peptide-centric immunopeptidomics, the HLA proteins from immunoprecipitation were also analyzed by LC-MS/MS. Combined with glycosylation analysis of both HLA proteins from both immunoprecipitation and flow-through, significant difference in glycosylation from HLA protein was found between control and infected cells. This find suggested that HLA synthesis and degradation was affected by SARS-Cov-2 infection, which led to the disruption of MHC-peptide complex formation and presentation of viral antigens.

## Results

### Immunopeptidomics Identified Viral Epitopes Presented on Both MHC I and MHC II

We designed an integrated immunopeptidomics and proteomics approach to characterize the interaction between SARS-Cov-2 and host cells. Human lung epithelial cell line Calu-3 was infected by the novel coronavirus at MOI of 5 for 24 hours. Then both the MHC class I and II immunopeptidome from control and infected cells were captured by immunoprecipitation with W6/32 and L243 conjugated agarose beads, respectively. Eluents from immunoprecipitation were further desalted and fractioned to keep immunopeptides and HLA proteins separately. The flow-through from immunoprecipitation were collected for proteomics analysis (Figure 1, Scheme for the workflow). Purified immunopeptides were analyzed by LC-MS/MS and immunopeptidome data were searched against a combined protein database from SARS-Cov-2 and human from Uniprot. The search results were filtered by applying 1% FDR to all the PSMs from each sample. A total number of 2695 and 2699 class I unique peptides were identified from control and infected cells, respectively while for class II peptides, 2632 and 2473 unique peptides were identified from control and infected cells, respectively. The accurate identification of immunopeptides was illustrated by the length distribution of both class I and class II peptides. For MHC I peptides, over 90% of identified peptides were found with 8-12 amino acids and the nearly 95 % of identified MHC II peptides had 13-25 amino acids (Figure S1). The immunopeptidome from the host between control and infected cells showed high level of overlap as similar percentage (60%) of MHC I and MHC II peptides were identified in both control or infected cells (Figure S2), which demonstrated the reproducibility in sample processing of the immunopeptidomcis workflow. Within the immunopeptidome, 3 Class I and 3 Class II peptides from SARS-Cov-2 were identified from infected cells and none of these peptides was identified from control experiments. The identification of viral epitopes was further validated by manual interpretation of MS/MS spectra (Figure S3). TGSNVFQTR from spike glycoprotein (638-646) was predicted binding to A*68:01 allele and it locates within the SD1 domain. ITFGGPSDSTGSNQNGER from nucelocapsid protein (15-32) was also identified in this study and has been predicted to bind to the same allele. Additionally, two class II peptides from nucleocapisd protein SPDDQIGYYRRATRRIR and FSKQLQQSMSSADSTQA were identified, which has not been reported before.

**Figure 1.**
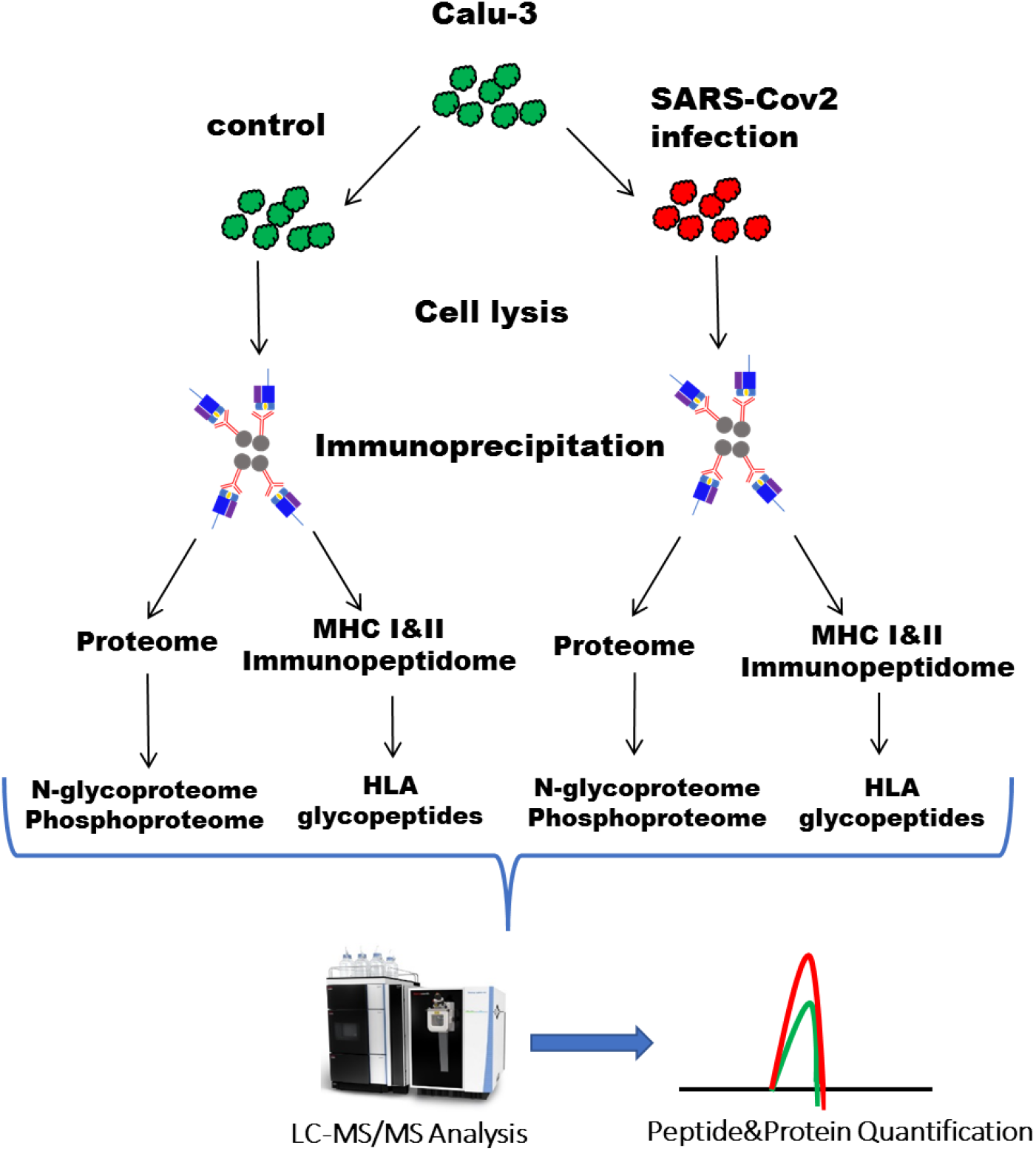
Scheme for the integrated immunopeptidomcis and proteomics analysis of SARS-Cov-2 infection of Calu-3 cells

### SARS-Cov-2 Infection Impacts Antigen Presentation and Activates Interferon Signaling Pathway

To further characterize the host response to viral infection and study the mechanism by which the infection affects the antigen presentation, additional proteomics studies were carried out. Following immunopeptidomics, protein contents from flow-through of immunoprecipitation were collected by acetone precipitation, followed by in-solution trypsin digestion. Protein digest from each sample was analyzed by LC-MS/MS with data-dependent acquisition and the relative abundance of protein was quantified by l*abel-free* quantification (LFQ) with Maxquant. Firstly, we evaluated reproducibility of the work-flow by plotting the LFQ intensities of identified proteins from both control and infection groups (Figure S4). These data showed a high correlation (R^2^>0.98, P<0.001 by linear regression) between biological replicates within each group, confirming that immunoprecipitation did not affect the proteomics analysis of the flow through. With an FDR of 1% at both peptide-spectrum match and protein level, a total number of 4479 proteins were identified, including 7 proteins from SARS-Cov-2 as shown in Figure 2. These proteins included ORF9b, nucleocapsid protein and spike glycoproteins, which has a LFQ intensity over 10^9^.

**Figure 2.**
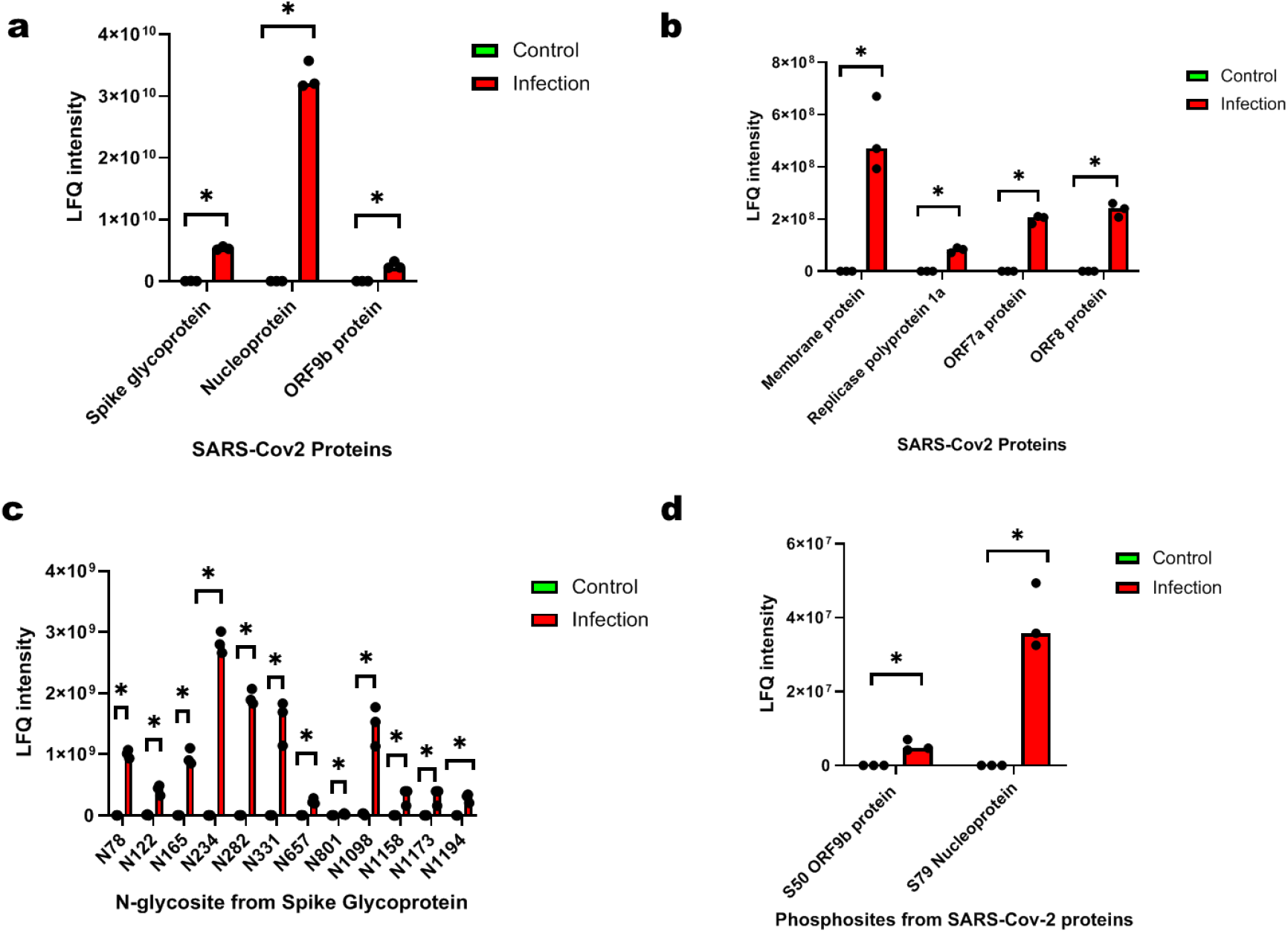
Abundance of viral proteins from control and infected Calu-3 cells. a, high abundant viral proteins; b, low abundant viral proteins; c, N-glycosites from spike glycoprotein; d, phosphosites from nucleoprotein and ORF9b.

Other viral proteins like membrane protein, replicase polyglycoproein 1a, ORF7a and ORF8 protein were found with lower abundance. Overall, 183 proteins were found with significant difference in abundance between control and infected cells by student’s t-test (fold change> 1.5, p<0.05) (Figure 3, a). The changes in host proteome can be attributed to the high expression of viral proteins as iBAQ quantification revealed that the most abundant viral protein, nucelocapsid protein ranked 18 in the all the identified proteins while spike glycoprotein ranked 481 (Figure 3, b). In order to analyze the functions of host proteins, gene ontology was applied to proteins with significant expression changes to better understand patterns of changes in cellular pathways or activities. These data showed that changes in protein abundance were predominantly enriched in antigen processing, transportation and presentation Go terms. These host protein pathways are noted to be involved in response to virus, including binding to virus and processing of viral proteins for antigen presentation (Figure 4, a). Of note, proteins participated in antigen processing (TAP1, TAP2 and TAPBP) were found with doubled expression in infected cells (Figure 4, b). Increased level of HLA-B and HLA-C (1.5 fold, p<0.05) were found in infected cells while no significant change in abundance of HLA-DR was found between control and infected cells (Figure 4, c). The HLAs in the proteome sample are from the flow-through of MHC immunoprecipitation, suggesting that the increased level of HLA-B and HLA-C represents HLAs that did not form peptide complexes. Reactome pathway analysis was used and identified pathways implicated in both innate and adaptive immune response to infection (Figure 5, a). This included interferon signalling pathway, an important part of innate immune system and pathway related to cytokine signaling, indicating the secretion of cytokines during infection. The activation of immune system also triggered pathways involving interferon-stimulate genes as 6 ISGs were found with significantly increased expression in infected cells (Figure 5, b). 3 members of STATs were identified with varied abundance change as STAT1 has an over 2 fold change in infected cells and STAT3 has only subtle change (1.2, p<0.05). Interestingly, no significant difference from STAT2 was found.

**Figure 3.**
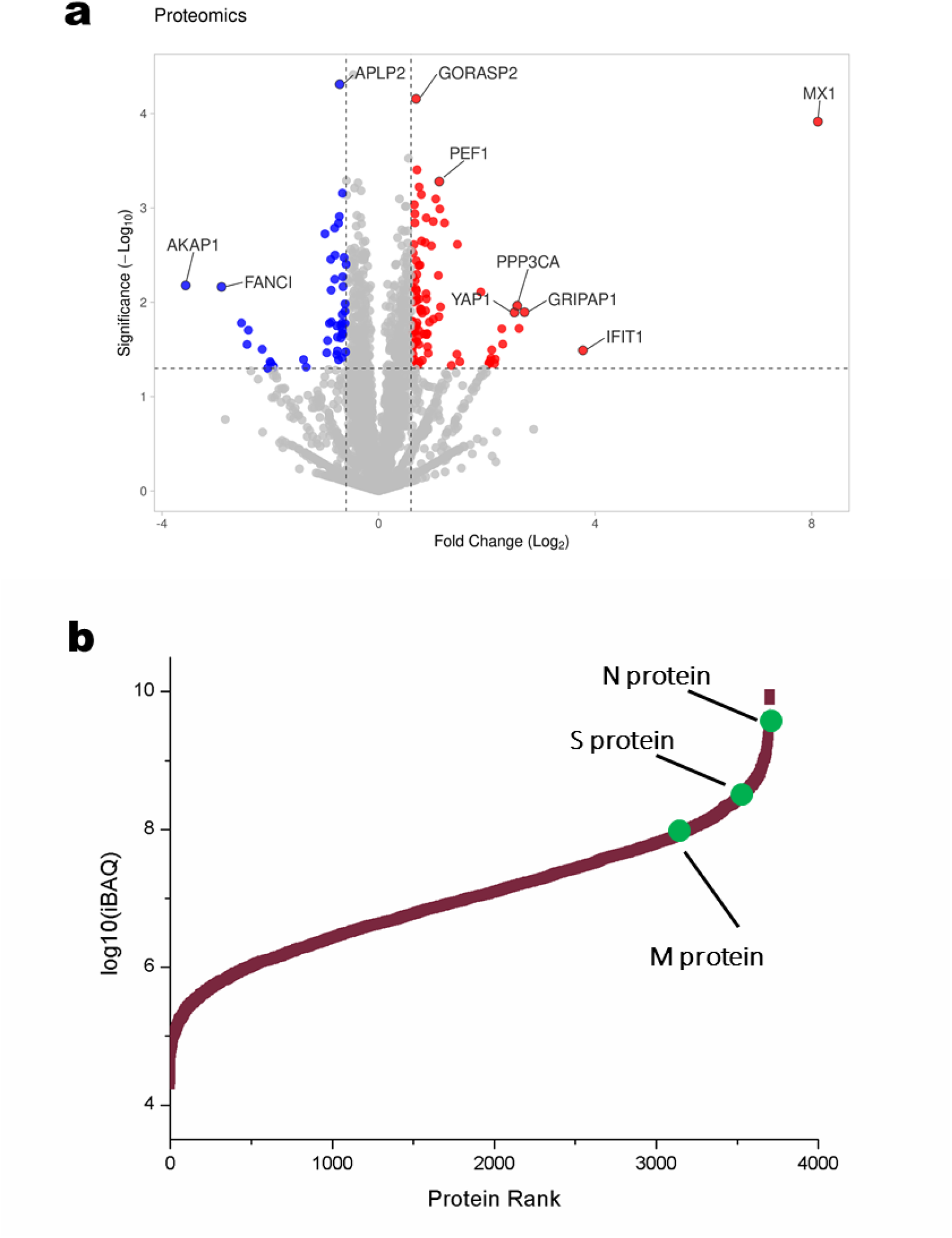
SARS-Cov-2 infection reshaped the proteome of Calu-3 cells. a, volcano plots of host proteins showing the difference in proteome; b, iBAQ quantification estimates the abundance of viral proteins in infected cells

**Figure 4.**
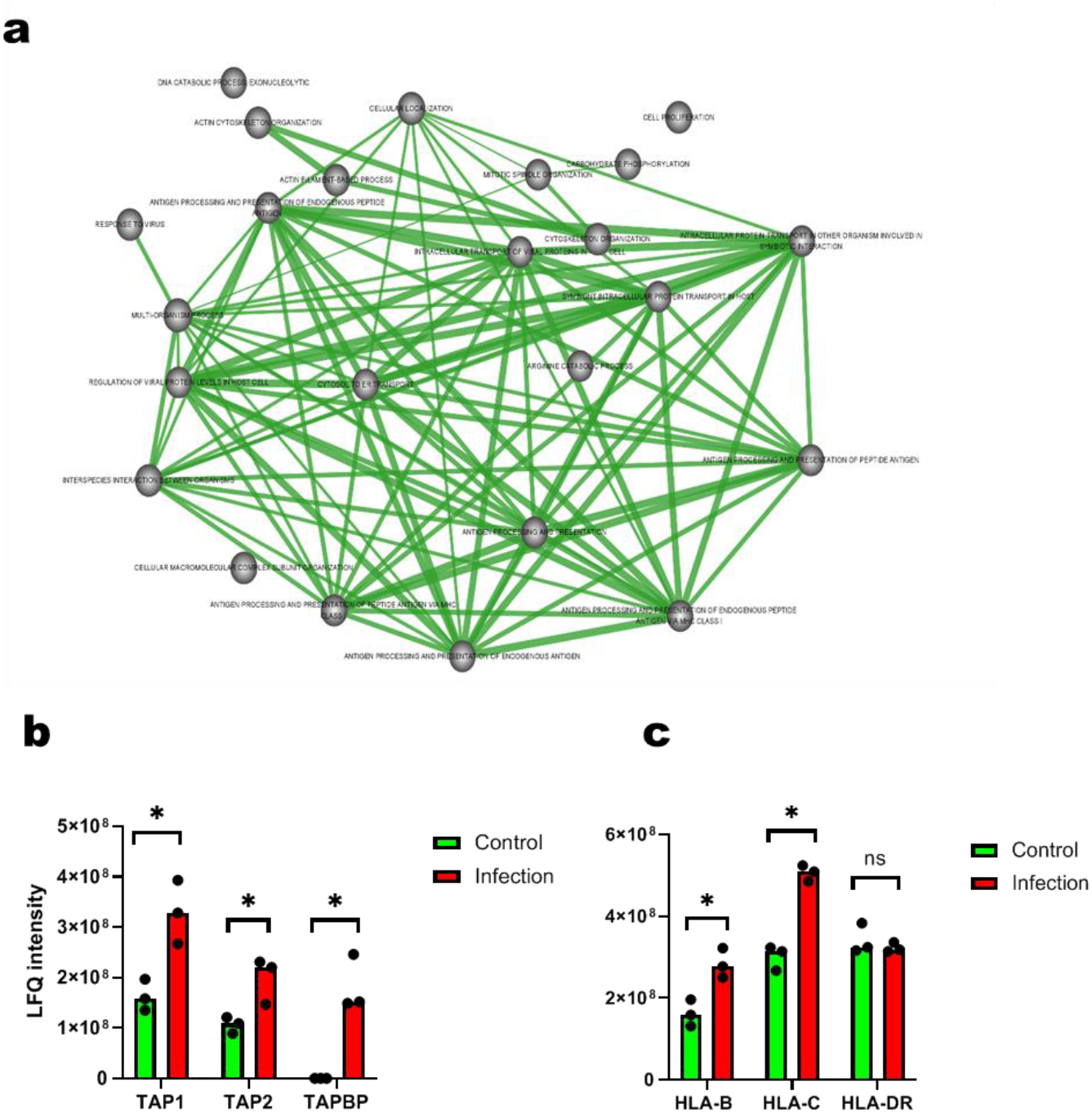
SARS-Cov-2 infection impacts antigen processing and presentation. a, Gene Ontology analysis showed enrichment in antigen and processing presentation from up-regulated proteins after infection; b, increased abundance of antigen processing proteins in infected cells; c, abundance of HLA-B and HLA-C are significantly higher in from the infected cells while there is no difference in the abundance of HLA-DR.

**Figure 5.**
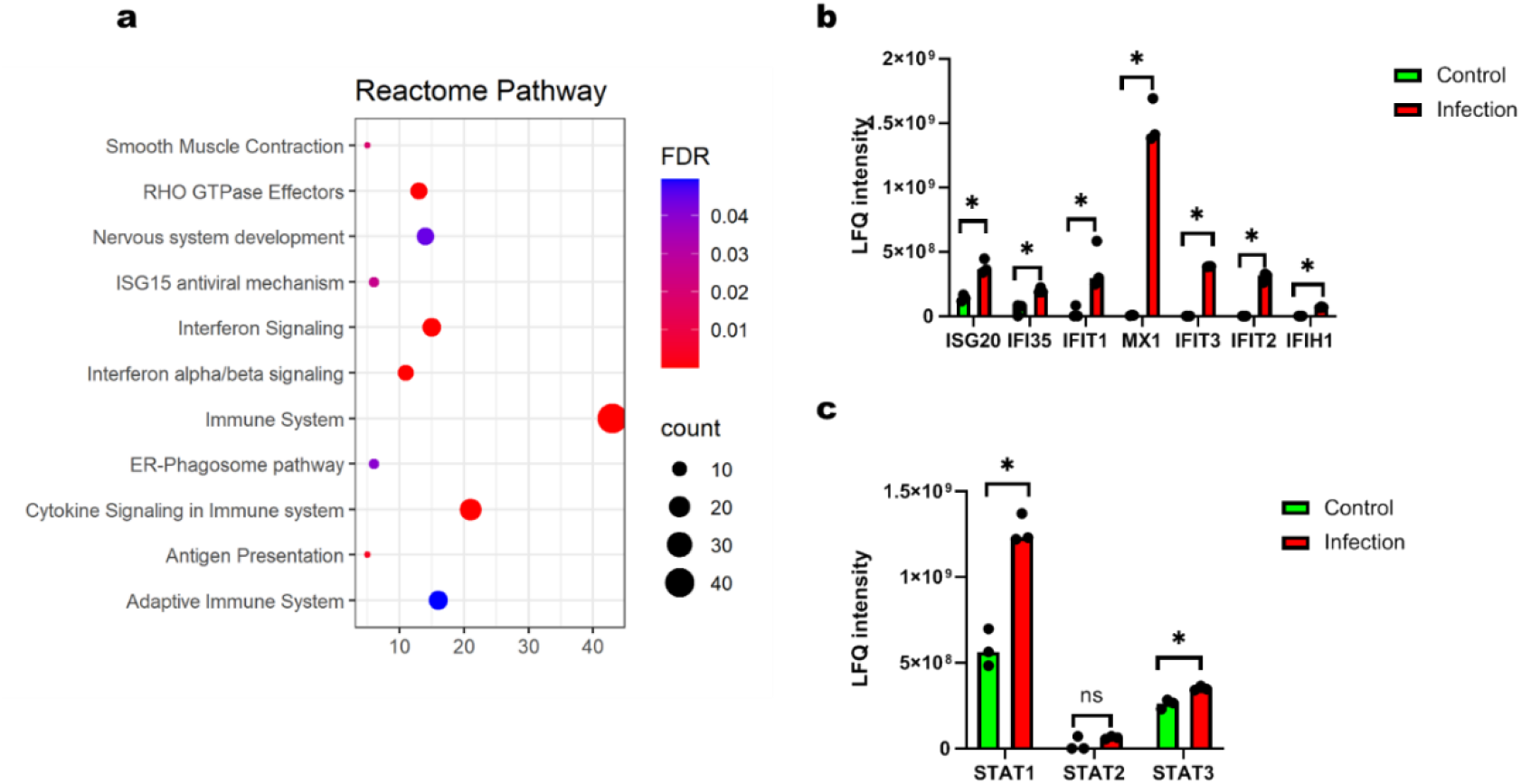
SARS-Cov-2 infection activates interferon signalling pathway. a, reactome pathway analysis reveals activation of type I interferon signaling pathways and immune response to virus infection and secretion of cytokines; b, increased expression of interferon stimulated genes in infected cells; c, increased expression of STATs in infected cells.

### Glycoproteomics Identifies Down-regulation of ACE2 and Increased Level of TLR3

It is well known that SARS-Cov-2 enters host cells through the binding of its spike glycoprotein to the ACE2 on the surface host cells^31^. Thus, it is also important to study the changes in the cell surface proteome to identify the receptors to sense the viral infection and trigger the down-streaming signaling. For this purpose, we applied quantitative glycoproteomics approach to quantify the N-glycosites from glycoproteins. Total 2142 N-glycosites with N#XS/T motif from 876 glycoproteins were identified and nearly all identified glycoproteins were annotated as cell surface and membrane proteins by Gene Ontology analysis (Figure S5). However, statistics demonstrated less change in the N-glycosites as only 91 N-glycosites showed significantly changes in infected cells compared to control cells (Figure 6, a) and those N-glycosites are listed in Supplementary File 3. Interestingly, ACE2 level in infected cells dropped by 2 fold compared with controlled cells through the quantification of N-glycosites containing Asn103 (Figure 6, b). Other glycoproteins involved in immune response to viral infection was also identified, including TLR3. Its Asn124 N-glycosite was significantly increased in infected cells with 5 fold increase in abundance and 7 other N-glycosites showed consistent trends (Figure 6, c, Figure S6), which demonstrated increased TLR3 expression in infected cells. Difference in glycosite level were also found with HLA proteins as up-regulated abundance of N-glycosite from HLA-B was found in infected cells (Figure 6, d). A non-classical MHC molecule HLA-E was also found with increased level from the proteome sample in infected cells (Figure 6, e) while no significant difference in the glycosites from HLA-DR(Supplementary File 3).

**Figure 6.**
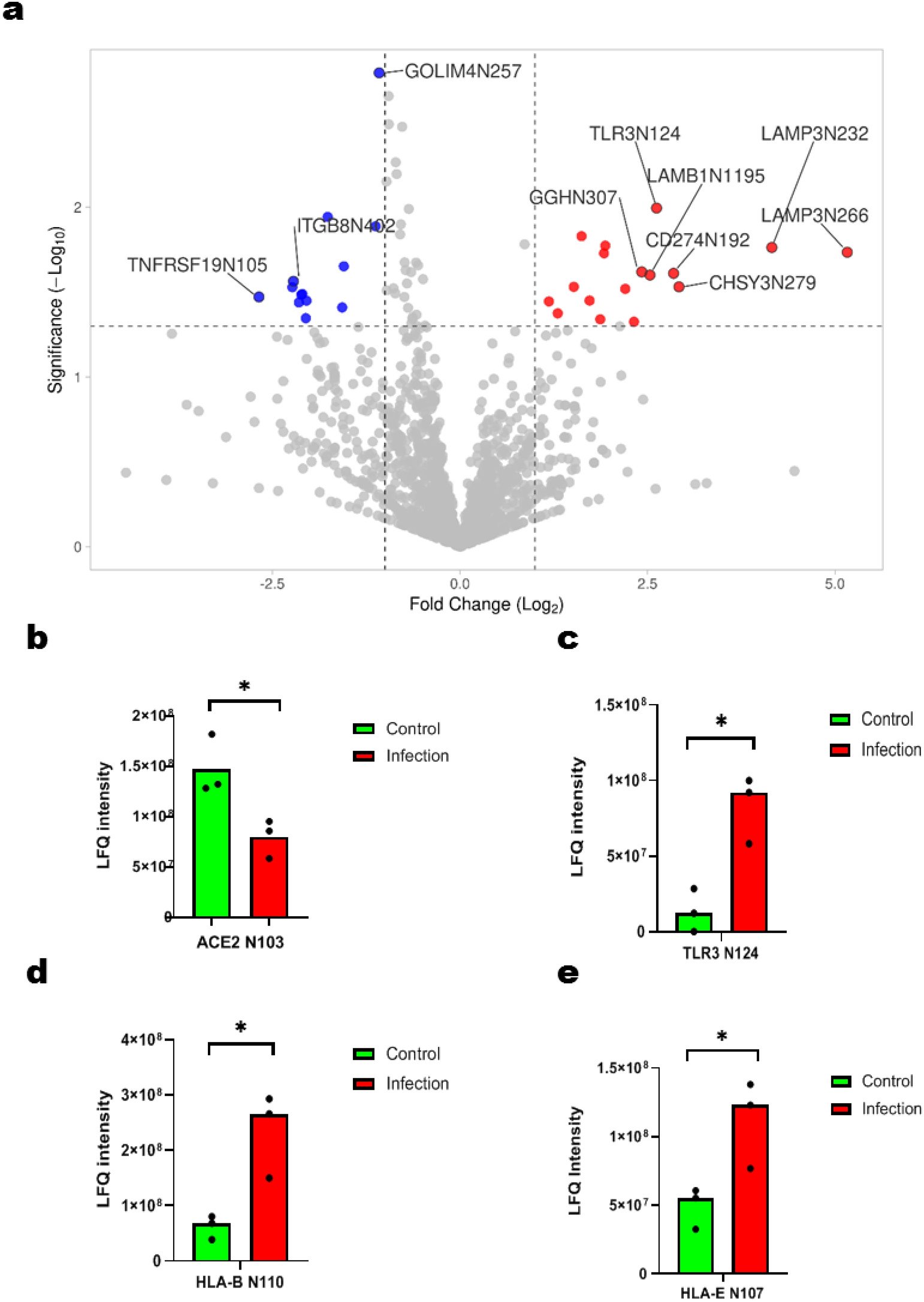
Glycoproteomics identified changes in the abundance of membrane proteins. volcano plots of N-glycosites quantified by glycoproteomics analysis (a); statistics of N-glycosites from proteins that involved in immune response to SARS-Cov-2 infection, including ACE2(b), TLR3(c),HLA-B(d) and HLA-E(e).

### Increased Level of Monoglucosylated and Truncated Glycosylation indicates HLA Degradation in Infected Cells

To confirm that the MHC complex was disrupted in infected cells, we expanded the glycoproteomics to study the glycosylation of HLA proteins from both immunoprecipitation and the flow-through lysate, which represent the HLAs that bound to peptides within MHC complex on the cell surface and “free” intracellular HLAs without forming complex. In conventional immunopeptidomics, after immunoprecipitation of MHC complex, only the peptides fraction was analyzed for the identification of epitopes binding to HLAs while the HLA proteins were discarded, which might lead to the loss of useful information on the structure and interaction of HLA proteins. In this study, the HLA protein fraction was collected by fractionation during C18 desalting, followed by digestion and LC-MS/MS. Database search showed high sequence coverage of HLA protein and it also identified proteins that interacts with MHCs (Supplementary File 4). We further analyzed the glycosylation of HLAs proteins from both the elution and flow-through of immunoprecipitation as the proper folding and antigen incorporation of HLA protein are dependent on the glycosylation^32,33^. Intact glycopeptides were enriched by HILIC-SPE and analyzed by LC-MS/MS directly. By database search with MSFraggerGlyco, the glycans attached to the HLA proteins were identified and it clearly showed difference in glycosylation from immunoprecipitation and the flow-through lysate (Figure 7, a). HLA proteins from the elution, which formed MHC-peptide complex, carried more hybrid and complex glycans that went through the complete glycan synthesis pathway^32,34^. On the other hand, HLA proteins from the flow-through lysate, which is “free” of complex and peptide, are found with mostly high-mannose type glycans, including the precursor with mono-glucosylation (Hex(10)HexNAc(2), Figure 7, b). Glycan analysis showed little difference between the immunoprecipitated HLA proteins from control and infected cells. However, from the flow-through lysate, higher level of glycopeptides (fold change ratio of 3.2) with mono-glucosylated in infected cells compared with controls (Figure 7, c) while glycopeptide with high mannose type glycan increased by 2 folds(Figure 7, d) and it is consistent with the fold change from the deglycosylated peptides. On the other hand, glycopeptides with fewer than 5 hexoses (Figure S7), which can be products of protein degradation, were only found in infected cells but not control cells (Supplementary File 6). By comparing the glycosylation pattern of HLA proteins, we can conclude that SARS-Cov-2 infection changed the synthesis and degradation of HLA proteins in Calu-3 cells and thus antigen presentation of MHC Class I system was disrupted.

**Figure 7.**
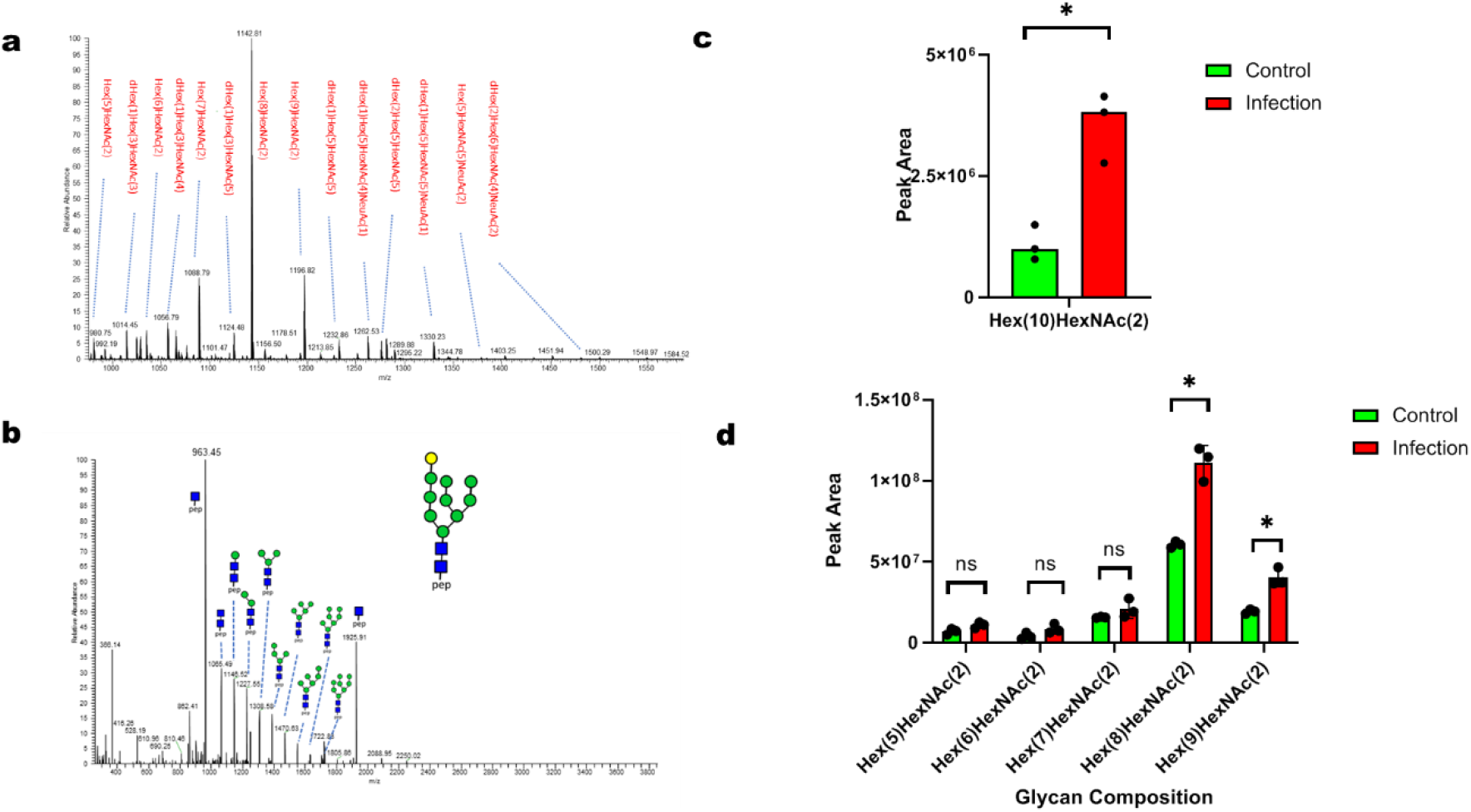
Glycosylation analysis of HLA glycosylation reveals possible HLA degradation after infection. a, glycan compositions identified from HLA protein captured by immunoprecipitation; b, MS/MS spectrum of mono-glucosylated HLA glycopeptide from infected Calu-3 cells; c, quantification of glycopeptides reveals increased level of mono-glucosylation of HLA after SARS-Cov-2 infection; d, statistics of normal high-mannose type glycans from HLAs showed no significantly difference between control and infected cells.

### Down-regulated AKT Signaling Identified by Phosphoproteomics

To further understand the host response to SARS-Cov-2 infection, we applied phosphoproteomics to elucidate mechanisms of cellular pathways and processes upon infection. Several viral proteins were found phosphorylated with various levels, including ORF9b, membrane protein and nucleoprotein. Infection also reshaped the host phosphoproteome as 181 phosphosites were found with significantly changed phosphorylation level (Figure 8, a). Pathway analysis found the reduced level of phosphorylation are enriched with proteins involved with AKT signaling(Figure 8, b), which was also found in a proteo-transcriptomomics of SARS-Cov-2 infected cells^35^. Phosphoproteomics also provided evidence in the changes in ErBB2 phosphorylation as varied levels in three sites were quantified with significant changes while the protein level remained the same (Figure 8, c, d). ErBB2 were also identified from protein fractions from MHC I and MHC II immunoprecipitation, suggesting it is interacting with MHC complex (Figure S8). However, it was not identified from the MHC immunoprecipitation with A549 and MDA 231 cells (Supplementary File 8). Thus, this interaction can be HLA allele dependent and requires further exploration.

**Figure 8.**
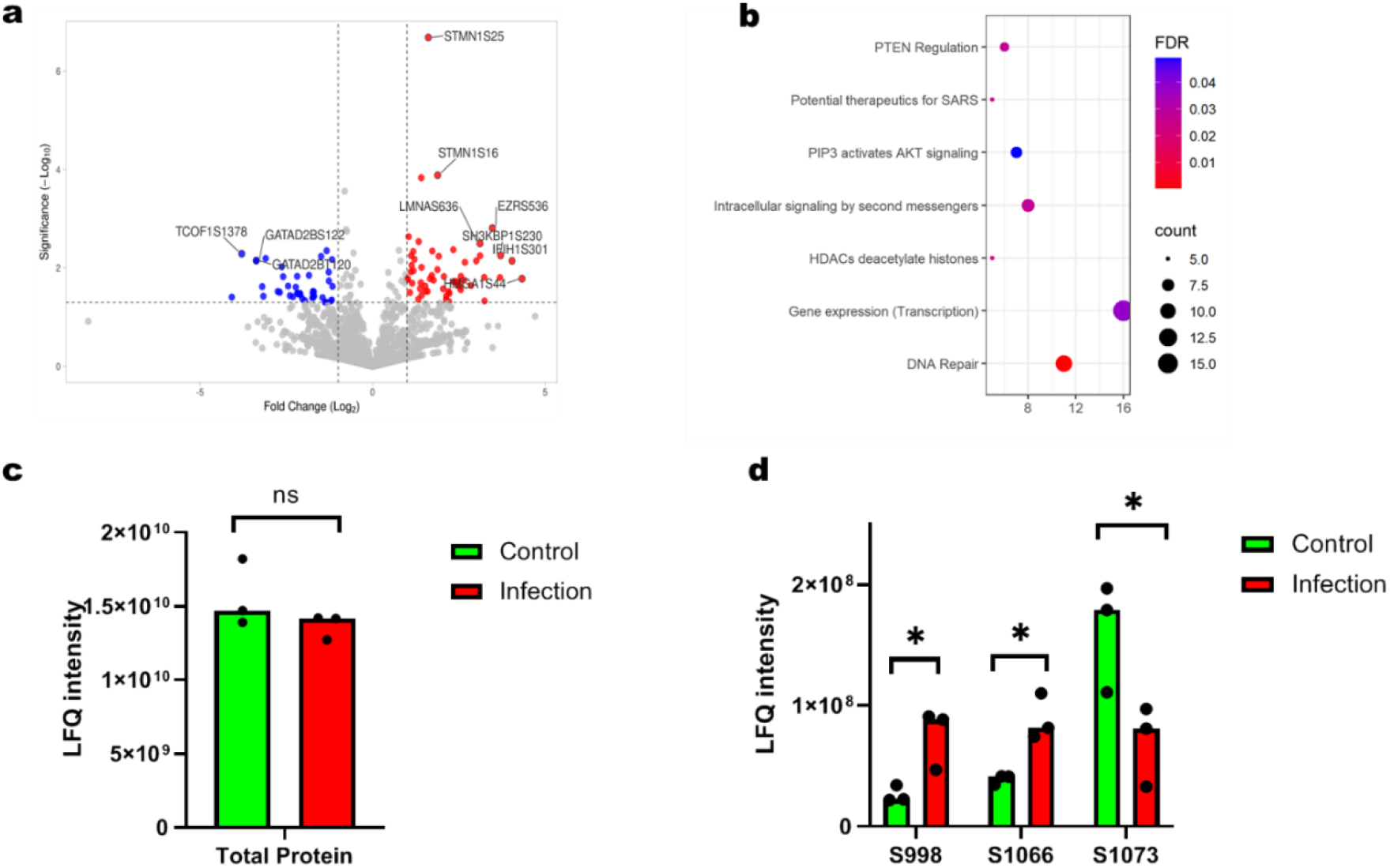
Phosphoproteomics identifies altered signalling in infected cells. a, volcano plots of phosphosites quantified by phosphoprteomics analysis; b, pathway analysis identified down-regulated pathways in infected cells; c, protein abundance of ErBB2 by total proteome analysis; d, different level of phosphorylation from multiple ErBB2 phosphosites identified by phosphoproteomics.

## Discussion

Host cells infected by the coronavirus is the simplest model to study the mechanism of viral infection and its impact on the immune system for the development of vaccines and therapeutics. Although several immunopeptidomics and proteomics studies of SARS-Cov-2 infected cells have been reported^19,23-26,36^, there is not yet a study that integrated both immunopeptidomics and proteomics, including PTM targeted sub-omics. Thus, we developed a workflow that included immunopeptidomcis, proteomics, glycoproteomics, phosphoproteomcis and HLA targeted interactomics to provide with a comprehensive overall view of how host cells response to infection that may attribute to the pathology of Covid-19.

With immunopeptidomics, epitope from spike protein was identified from MHC I immunopeptidome while epitopes from nucleocapsid protein was identified from both MHC I and MHC II. A few viral epitopes were also reported from previous study with Calu-3 cell lines and matched its HLA alleles. Additionally, this study provided the first identification of MHC II epitopes from nucleoprotein and one of the peptide (SPDDQIGYYRRATRRIR) has a few overlaps with a class I epitope that identified from a previous study (NSSPDDQIGYY) and this epitope was found immunogenic in eliciting CD^8+^ T cells. Further study is needed to investigate the immunogenicity of those viral epitopes and nucleopcapsid protein could also been considered as a target for vaccine development since its sequence has been stable even with the emergence of different variants of SARS-Cov-2. Fewer viral epitopes were identified in this study with Calu-3 cells compared with a recent study with HEK293 and A549 cell lines that transfected with ACE and TMSSP2^19^. We suspect that it could be related to the difference in HLA alleles among the cell lines as downregulation of surface MHC I was found in Calu-3 cells. To investigate how the antigen presentation was affected by viral infections, we performed total proteome analysis with the flow-through from MHC immunoprecipitation and higher level of HLA-B and HLA-C were found from infected cell while the level of HLA-DR remained the same. The result seems contradictory to recent discoveries that MHC-I was down-regulated in infected cells and Covid-19 patients^28,29^.However, since the proteomics samples are from the flow-through of MHC immunoprecipitation, it can be anticipated that the HLA proteins for proteomics analysis are actually free of MHC complex and the increased abundance of HLA-B and HLA-C are more likely from disrupted MHC complex. This is consistent with the finding the MHC-I molecules are selectively targeted for lysosomal degradation via autophagy by ORF8, which was found in infected cells^28^. To prove that HLA was indeed degraded before forming complex with antigens and beta-2-micorgloblin, we performed glycosylation analysis of HLAs from both the elution and flow-through of MHC immunoprecipitation. It was revealed that HLA proteins have different glycosylation when it was associated with antigens on the cell surface compared when it was antigen-free. Higher level of mono-glucosylated HLA glycopeptide indicated that certain amount of HLA proteins was hindered from proper folding and resulted in the gradation of HLA protein and generation of trimmed glycopeptides with less than 4 mannoses. Higher level of improperly glycosylated HLAs in infected cells suggested that SARS-Cov-2 infection changed the folding and degradation of HLA proteins and more HLAs were stopped forming MHC-complex. However, this change is too subtle to be detected by either total proteomics or immunopeptidomics alone as suggested from previous studies. Interactomics of viral proteins proved that several viral proteins interacts with HLAs in human cell lines^21,37^, however, those proteins were not identified in the digest of HLA proteins from MHC immunoprecipitation, suggesting they only bind to HLA proteins intracellularly.

Thus, it is important to include multiple level of –omics and analyze the data systematically. With the addition of quantitative glycoproteomics, we discovered the strong innate immune response in infected cells could be initiated by TLR3, which senses viral RNA and triggers the activation of interferon signalling pathway. The increased expression of TLR3 suggested that the activation of interferon pathway was triggered by recognition of dsRNA from SARS-Cov-2, followed by secretion of cytokines. A recent study suggested the stimulation of INF pathway in SARS-Cov-2 infected Calu-3 cells was due to the lower expression of M protein. However, M protein also has a high expression in Calu-3 cells after 24 hours. We hypothesize that activation of interferon pathway can be attributed to the higher expression of TLR3 in Calu-3 cells, which has higher transcription of TLR3 gene compared to Caco-2 and A549 cell lines in other proteomics studies ^38^. This assumption is partly supported by the find that inhibition of histamine-induced expression of TLR3 in SARS-CoV-2 infected cells can reduce TLR3-dependent signaling processes and secretion of cytokines^39^. This is the first study that links TLR3 with SARS-CoV-2 sensing specifically and TLR3 is well known to activate the production of inflammatory cytokines and triggers expression of interferon-inducible genes through the JAK-STAT pathway^40^. Among the three STAT members identified, STAT1 showed highest fold change and it is consistent with previous find that STAT1contributes to innate immune response during viral infection ^41^. Proteomics identification also suggest potential roles of ISGs in antiviral activity, including ISG20, which has exonuclease activity with specificity for single-stranded RNA. The antiviral activity of IFN against RNA virus is partly mediated by ISG20^42^. It also discriminates between self and non-self translation ^43^. Thus, we can conclude that SARS-Cov-2 infection also triggered the innate immune response in Calu-3 cells through TLR3-STAT1-INF-ISG axis.

With this integrated study, we revealed that there is imbalanced innate and adaptive immune response in infected Calu-3 cells, which was not discovered in other similar study with different cell lines. Thus, we suspect that the impact of viral infection on antigen presentation could be HLA allele dependent as the association between HLA allele frequencies and susceptibility to Covid-19 in large population was suggested by a few studies^44,45^. Interestingly, one HLA allele of Calu-3 cells, HLA-B*51:01, has been found in association with severe disease and worse outcomes in Chinese population with Covid-19^46^. The impaired antigen presentation with these HLA alleles may lead to the exhaustion of CD^8+^ T cells in patients with severe symptoms^30^ while CD^4+^ T cell response can be found in most patients and unexposed individuals ^8^, which is consistent with our finding that the presentation of MHC II antigens are not affected in infected cells. In conclusion, we conducted an integrated immunopeptidomics and proteomics study of Calu-3 cells infected by SARS-Cov-2. It was revealed that while innate immune system was activated through TLR3-STAT1-INF-ISGs axis, the adaptive immune system was affected even though viral antigens were identified from both MHC I and MHC II immunopeptidome. The mechanism of down-regulation of MHC I antigen presentation was further studied by comprehensive analysis of HLA glycosylation as higher level of improper glycosylation was found in infected cells ant it led to the degradation. Expanded application of the established approach to more cell lines with difference variants and clinical samples with accurate HLA genotyping can shed more lights on how impaired antigen presentation contribute to the pathology of Covid-19 and help develop precision strategy for vaccination and therapy.

## Materials and Methods

### Cell Culture and Infection

Calu-3 cells were cultured at 37 °C and in 5 % CO_2_ in DMEM medium supplemented with 10 % fetal bovine serum (FBS), 50 IU/mL penicillin, 50 μg/mL streptomycin and 2 mM L-glutamine. Cells are allowed to grow to 80-90 % confluence in T175 flasks. Vero and Vero E6 cells were cultured in DMEM supplemented with 1X non-essential amino acid, 20 U/mL Penicillin, 0.02 mg/mL Streptomycin, 1 mM sodium pyruvate and 5% FBS. SARS-CoV-2 isolate Canada/ON/VIDO-01/2020 was propagated on Vero E6 cells and titered on Vero cells. Exact genetic identity to original isolate was confirmed by whole viral genome sequencing. Passage three virus stocks were used in all subsequent experiment that required live virus. For cell infection, cells were washed with PBS and infected by SARS-Cov-2 at a MOI of 5 at 37 °C for 1 h in Infection media (DMEM supplemented with 1X non-essential amino acid, 20 U/mL Penicillin, 0.02 mg/mL Streptomycin, 1 mM sodium pyruvate). After infection, the cells were washed with Infection media and then cultured in 30 mL Infection media for a further 24 h at 37 °C. For controls, only media were changed and cells were kept for 24 h.

### Immunoprecipitation

Both control cells and infected Calu-3 cells were washed with 10 mL PBS 3 times to remove the media. 5 ml of lysis buffer (20 mM Tris-HCl, pH=7.4, 100 mM NaCl, 5 mM MgCl2, 0.5% CHAPS, 1.5% Triton X-100) with protease and phosphatase inhibitor was added to each flask. Cells were lysed by shaking the flask. Detached cells were transferred into 50 mL tube and rotated for 1h at 4 °C. Cell lysate was cleared by centrifugation at 16,000 × *g* for 10 min and supernatant were transferred into new tube for immunoprecipitation.

W5/32 (MHC I) and L243 (MHC II) antibodies were conjugated to Amino-link plus agarose beads following the manufacture’s instruction and 1 ml of resins were packed into a cartridge for immunoprecipitation. Cell lysates were firstly passed through an MHC I affinity column and the flow-through was immediately loaded onto MHC II affinity column. Flow-through from each immunoprecipitation was collected for proteomics analysis. Both MHC I and MHC II affinity columns were washed with 10 volumes of wash buffers as described previously ^47^ and MHC-peptides complex was eluted with 5 ml of 10 % acetic acid. MHC associated peptides were purified with C18 cartridge and eluted with 30 % acetonitrile. Eluted peptides were lyophilized by Speed Vac and reconstituted in 0. 1% formic acid for LC-MS/MS analysis. Remaining proteins fractions from immunoprecipitation were further eluted from the C18 cartridge by 80 % acetonitrile and lyophilized by Speed Vac. Dried protein was reconstituted in in 8 M urea and 100 mM Tris-HCl (pH=7.4), reduced by DTT, alkylated by IAA and digested by trypsin. Protein digest was desalted by HyperSep C18 SPE cartridge and protein digest were dried by Speed Vac for further process.

### Sample preparation for proteomics analysis

Cell lysate from the flow-through of immunoprecipitation was collected for proteomics analysis. Proteins from the flow-though were precipitated by adding 10 volumes of cold acetone overnight at -20 °C. Protein precipitates were washed with acetone for three times and air dried in a fume hood. For digestion, proteins were solubilized in 8 M urea and 100 mM Tris-HCl (pH=7.4) and protein concentration was measured by Dc Assay. Proteins were reduced by DTT, alkylated by IAA and digested by trypsin. Protein digest was desalted by HyperSep C18 SPE cartridge and protein digest were dried by Speed Vac for further process.

Hydrophilic interaction solid phase extraction (HILIC SPE) was conducted as previously described with minor changes^48^. 200 μg of dried protein digest was solubilized in 80 % of acetonitrile (ACN) with 1 % of trifluoroacetic acid (TFA) and loaded onto the microcolumn packed with 5 mg of polyhdroxylethyl beads (100 Å, 5 μm). After three washes with the same loading buffer, glycopeptides retained on SPE column were eluted by 30 % of ACN with 0.1 % TFA. For the flow-through lysate digestion, two replicates of HILIC enrichment were carried out and eluted glycoppeptides from one replicate was lyophilized for intact glycopeptides analysis. Glycopeptides from the other replicate was then deglycosylated by incubation with 50 μL of 50 mM Tris-HCl (pH=7.5) with 1 unit per microliter of PGNase F (New England Biolabs) over night at 37 °C. Deglycosylated peptides were dried with Speed Vac and the samples were stored at -80 °C until MS analysis. For HLA protein from immunoprecipitation, flow-though and wash was combined to collect non-glycosylated peptides for protein identification. Elution from HILIC SPE was lyophilized for intact glycopeptides analysis.

For phosphopeptide enrichment, affinity enrichment with titanium dioxide nanoparticles was adapted from a previous method^49^. TiO_2_ nanoparticles were prepared in suspension of 20 mg/ml in 0.1% TFA, 80% acetonitrile with the adding of glycolic acid (300 mg/ml). 200 μg protein digest was reconstituted in 200 μl of each loading buffer and 50 μl of TiO_2_ suspension was added. The mixture was kept shaking at room temperature for 30 min followed by centrifugation at 10,000 *g* for 3 min. Supernatant was discarded and 250 μl of loading buffer was added. The mixture was shaken for 15 min and centrifuged at 10,000 *g* for 3 min. The pellet was then washed by 200 μl of 80% ACN in 0.1 % TFA for 15 min. After centrifugation at 10,000 g for 3 min, 200 μl of 5% ammonia was added to the pellet and the pellet was shaken for 15 min. In the final step, the mixture was centrifuged at 17,000 *g* for 3 min and supernatant containing eluted phosphopeptides was transferred to a clean tube. Eluted phosphopeptides was dried the elution with Speed Vac and stored the samples at -80 °C until MS analysis

### LC-MS/MS Analysis

All the samples were analyzed by an LC-MS/MS system comprised of Exploris 480 mass spectrometer (Thermo Fisher Scientific Inc, Bremen, Germany) and Ultimate 3000 UHPLC. Peptides was were loaded via an Acclaim PepMap 100 trap column (C18, 5 μm, 100Å; Thermo Scientific) onto a Waters BEH C18 column (100 μm × 100 mm) packed with reverse phase beads (1.7 μm, 120-Å pore size, Waters, Milford, MA). A 15, 45 minutes or 90 minutes gradient from 5–30 % acetonitrile (v/v) containing 0.1 % formic acid (v/v) was performed at an eluent flow rate of 500 nL/min. For data-dependent acquisition, a full ms1 scan (resolution: 60, 000; AGC target: standard; maximum IT: 50 ms; scan range: 375-1600 m/z) preceded by subsequent ms2 scans (resolution: 15,000; AGC target: standard; maximum IT: 150 ms; isolation window: 2 m/z; scan range: 200-2000 m/z; NCE: 30). To minimize repeated sequencing 1 of the same peptides, the dynamic exclusion was set to 60 sec and the ‘exclude isotopes’ option was activated.

### Database Search and Analysis

For immunopeptidomics, database search was performed by MSfragger 3. 0 ^50^, Maxquant ^51^ and Peaks DB^52^ against Uniprot human protein database with common contaminants proteins, plus the protein sequences from SARS-Cov-2. Both precursor and fragments tolerance were set at 20 ppm. Non-specific cleavage was set and peptides with 7-25 amino acids were retained. All PSMs were filtered with a 1% FDR and the identified peptides were further filtered by peptide length (8-14 for MHC I peptides and 12-25 for MHC II peptides). Peptides from HLA and beta-2 microglobulin were also removed.

Data from intact glycopeptides analysis were processed by MSfragger using N-glyco-HCD workflow and the glycan composition was verified manually. For proteomics data, RAW data were processed with Maxquant ^51^ and protein database search were done by Androma engine^53^ against the same database used for immunopeptidomics. For total proteomics, carbamidomethylation of cysteine was set as fixed modification. Methionine oxidation and as variable modifications. For phosphopteomics, serine/threonine/tyrosine phosphorylation were added as variable modification. For N-glycoproteomics, deamination of asparagine was set as variable modification. An FDR of 1% was set for PSM, site and protein. LFQ was selected to extract the peak intensities of each phosphosites and the results were processed by Persus. Paired student T test was used to compare the abundance of proteins, phosphosites and glycosites. Functional analysis and the interaction network was analyzed by String^54^ and the results were visualized by Enrichment Map^55^ within the Cytoscope environment ^56^. Statistical analysis was performed with GraphPad Prism 9. Volcano plots were made by VolcaNoseR^57^.

## Data availability

The mass spectrometry proteomics data have been deposited to the ProteomeXchange Consortium via the PRIDE^58^ partner repository with the dataset identifier PXD036022.

## Acknowledgement

The authors thank the Collaborative Unit in Translational Research between National Research Council Canada and CHU Ste. Justine for funding support.

## Author contributions

R.C., K.M.F., S.M.T. and J-J.L. designed the omics research and wrote the manuscript. A.T and D.D designed and performed virus infection work. R.C. performed sample and data analysis for omics research. K.K and E.C. analyzed the immunopeptidomics data.

