## Supplementary material for "Integrated Immunopeptidomics and Proteomics Study Reveals Imbalanced Innate and Adaptive Immune Responses to SARS-Cov-2 Infection": Supplementary Figures.pptx

### Slide 1
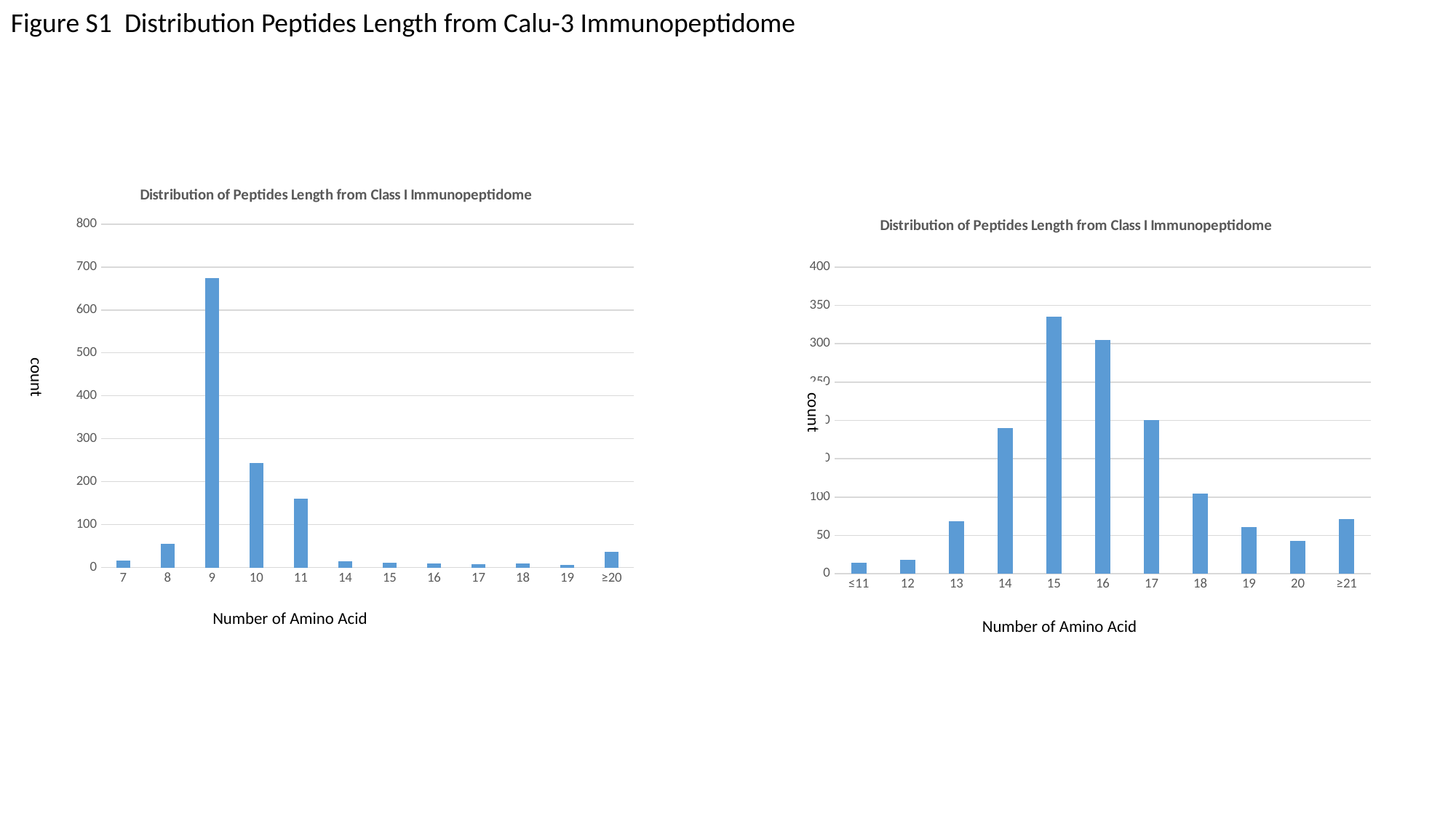

Figure S1 Distribution Peptides Length from Calu-3 Immunopeptidome
#### Chart: Distribution of Peptides Length from Class I Immunopeptidome
| Category | Count |
|---|---|
| 7 | 17.0 |
| 8 | 56.0 |
| 9 | 675.0 |
| 10 | 244.0 |
| 11 | 160.0 |
| 14 | 14.0 |
| 15 | 11.0 |
| 16 | 9.0 |
| 17 | 7.0 |
| 18 | 9.0 |
| 19 | 6.0 |
| ≥20 | 37.0 |
#### Chart: Distribution of Peptides Length from Class I Immunopeptidome
| Category | Count |
|---|---|
| ≤11 | 14.0 |
| 12 | 18.0 |
| 13 | 68.0 |
| 14 | 190.0 |
| 15 | 335.0 |
| 16 | 305.0 |
| 17 | 200.0 |
| 18 | 104.0 |
| 19 | 61.0 |
| 20 | 43.0 |
| ≥21 | 71.0 |count
Number of Amino Acid
Number of Amino Acid

### Slide 2
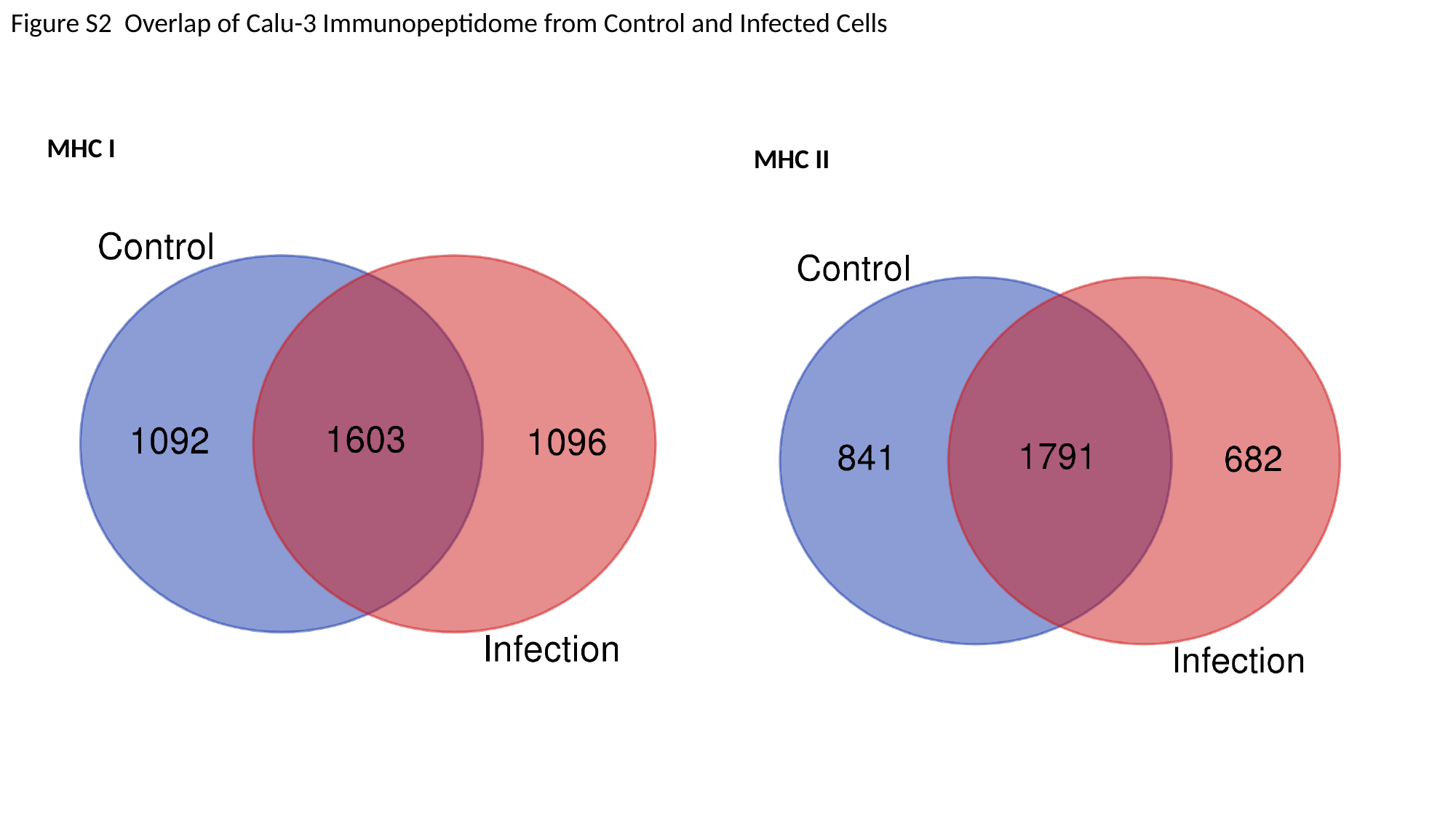

Figure S2 Overlap of Calu-3 Immunopeptidome from Control and Infected Cells
MHC I
MHC II

### Slide 3
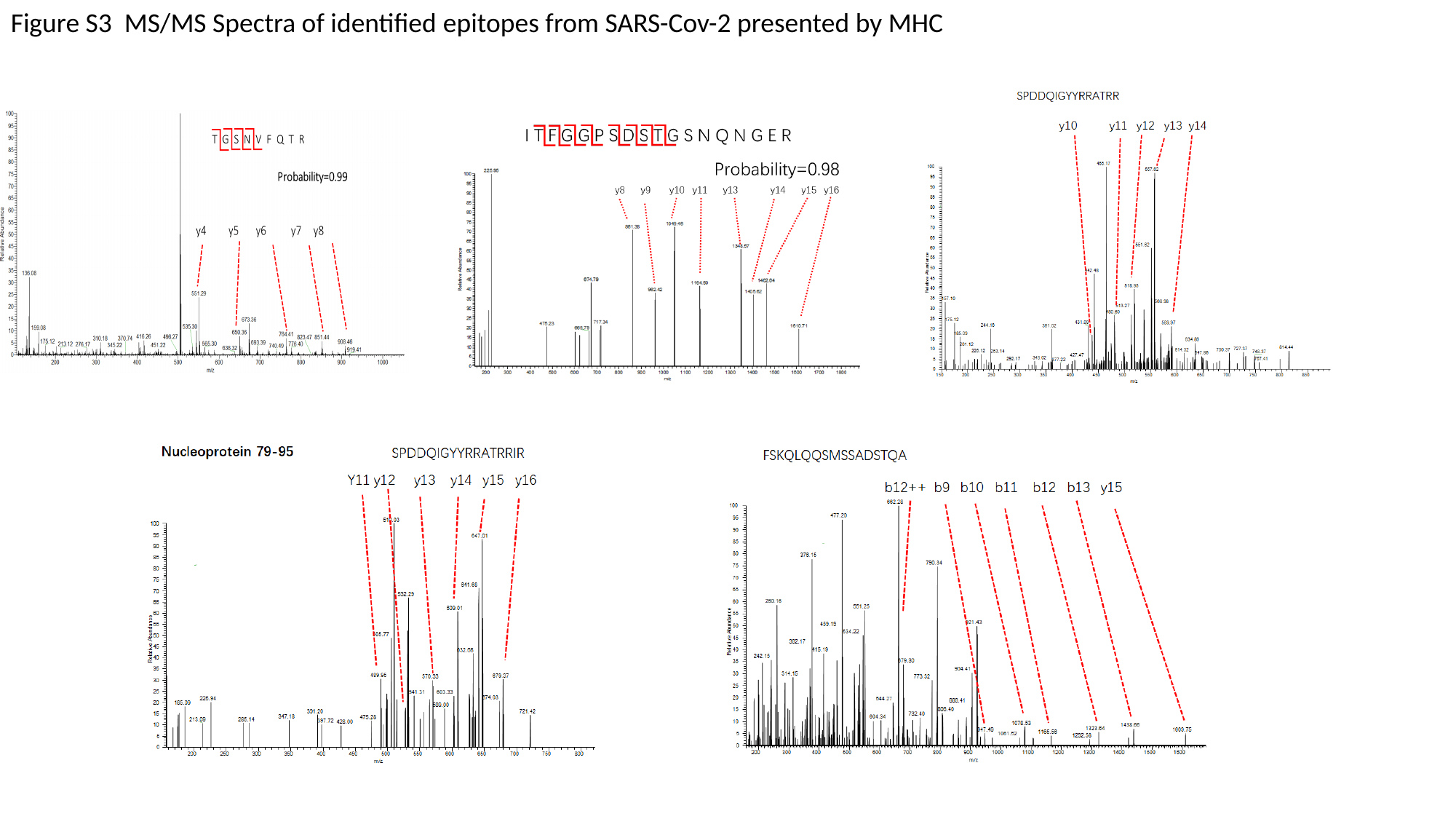

Figure S3 MS/MS Spectra of identified epitopes from SARS-Cov-2 presented by MHC

### Slide 4
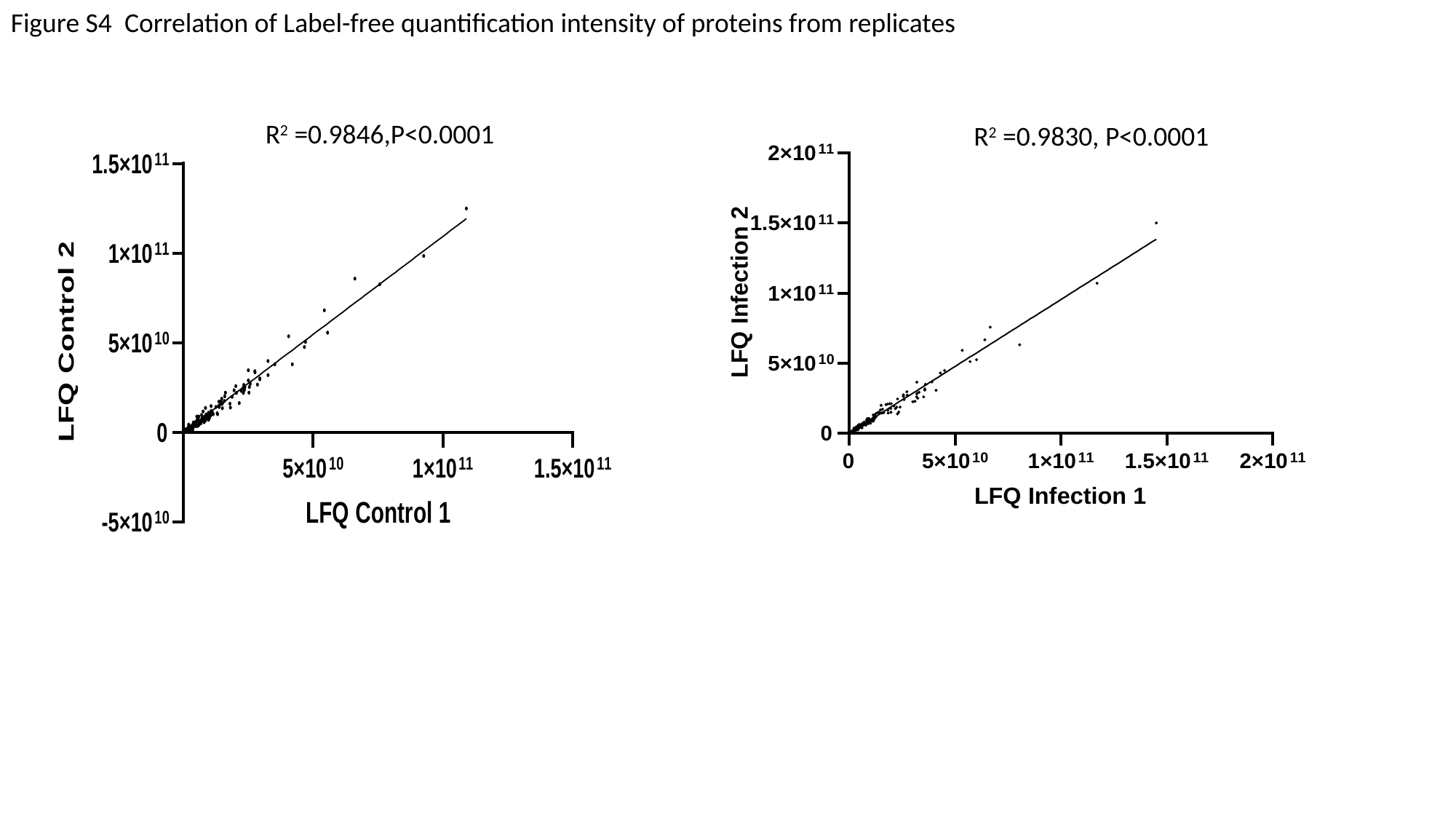

Figure S4 Correlation of Label-free quantification intensity of proteins from replicates
R^2^ =0.9846,P<0.0001
R^2^ =0.9830, P<0.0001

### Slide 5
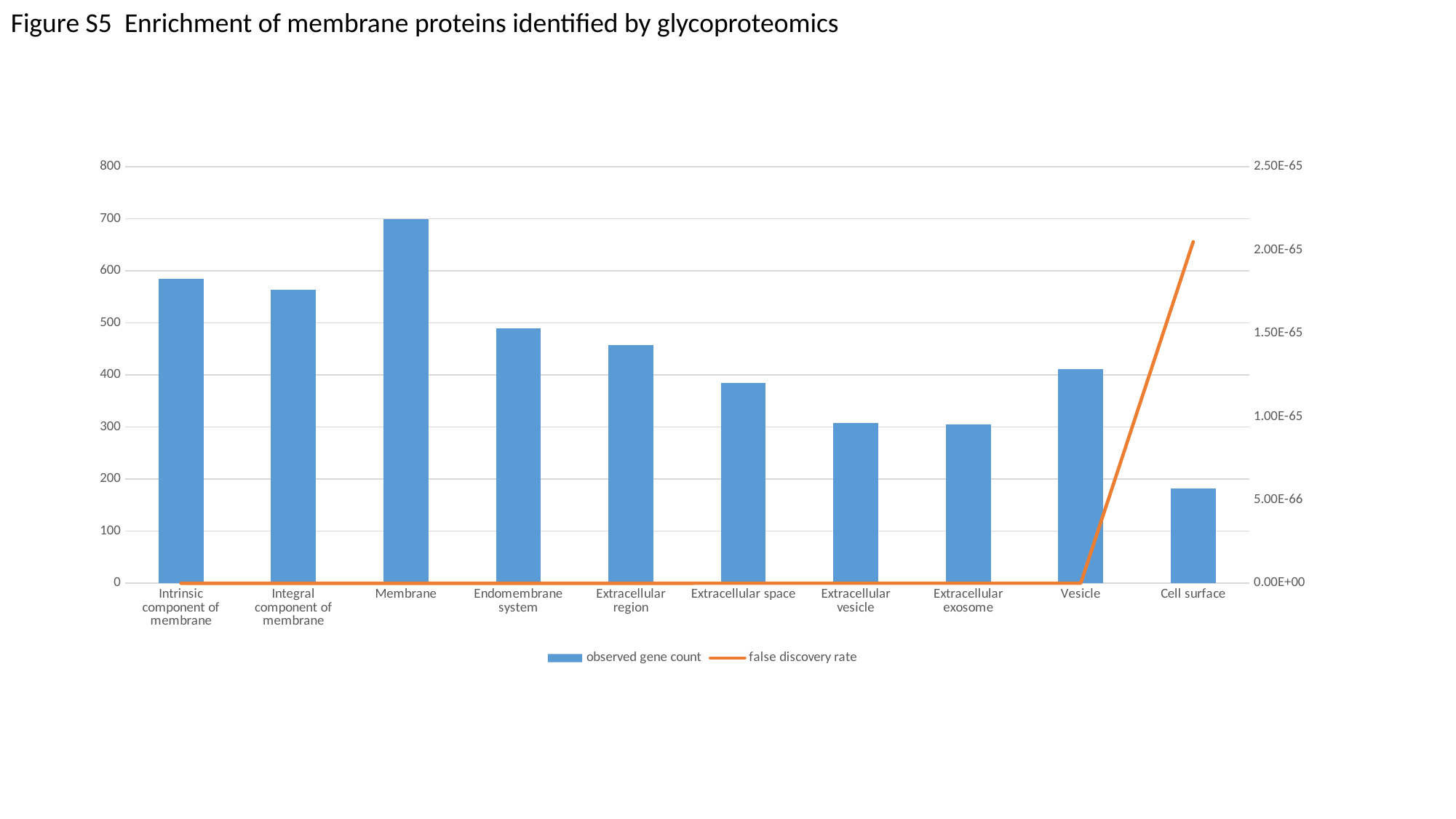

Figure S5 Enrichment of membrane proteins identified by glycoproteomics
#### Chart
| Category | observed gene count | false discovery rate |
|---|---|---|
| Intrinsic component of membrane | 585.0 | 3.61e-129 |
| Integral component of membrane | 564.0 | 1.81e-120 |
| Membrane | 699.0 | 3.26e-96 |
| Endomembrane system | 490.0 | 1.28e-94 |
| Extracellular region | 458.0 | 4.06e-88 |
| Extracellular space | 385.0 | 1.67e-79 |
| Extracellular vesicle | 308.0 | 5.58e-77 |
| Extracellular exosome | 305.0 | 3.19e-76 |
| Vesicle | 412.0 | 1.58e-71 |
| Cell surface | 182.0 | 2.05e-65 |

### Slide 6
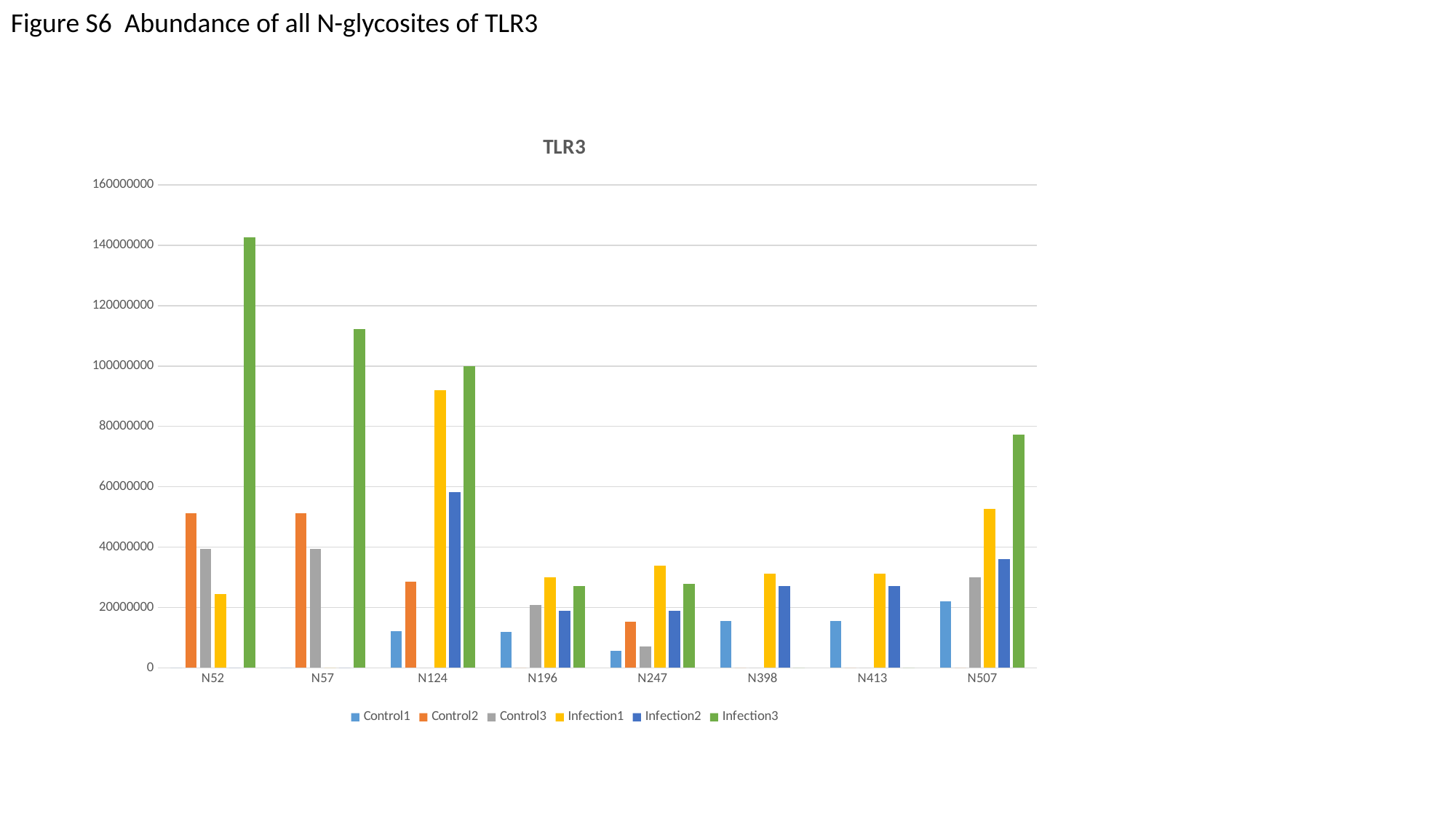

Figure S6 Abundance of all N-glycosites of TLR3
#### Chart: TLR3
| Category | Control1 | Control2 | Control3 | Infection1 | Infection2 | Infection3 |
|---|---|---|---|---|---|---|
| N52 | 0.0 | 51201000.0 | 39290000.0 | 24508000.0 | 0.0 | 142650000.0 |
| N57 | 0.0 | 51201000.0 | 39290000.0 | 0.0 | 0.0 | 112160000.0 |
| N124 | 12248000.0 | 28444000.0 | 0.0 | 92080000.0 | 58225000.0 | 99939000.0 |
| N196 | 11839000.0 | 0.0 | 20820000.0 | 30107000.0 | 18933000.0 | 27086000.0 |
| N247 | 5678800.0 | 15387000.0 | 7058500.0 | 33807000.0 | 18978000.0 | 27752000.0 |
| N398 | 15624000.0 | 0.0 | 0.0 | 31145000.0 | 27148000.0 | 0.0 |
| N413 | 15624000.0 | 0.0 | 0.0 | 31145000.0 | 27148000.0 | 0.0 |
| N507 | 22070000.0 | 0.0 | 30025000.0 | 52758000.0 | 36113000.0 | 77258000.0 |

### Slide 7
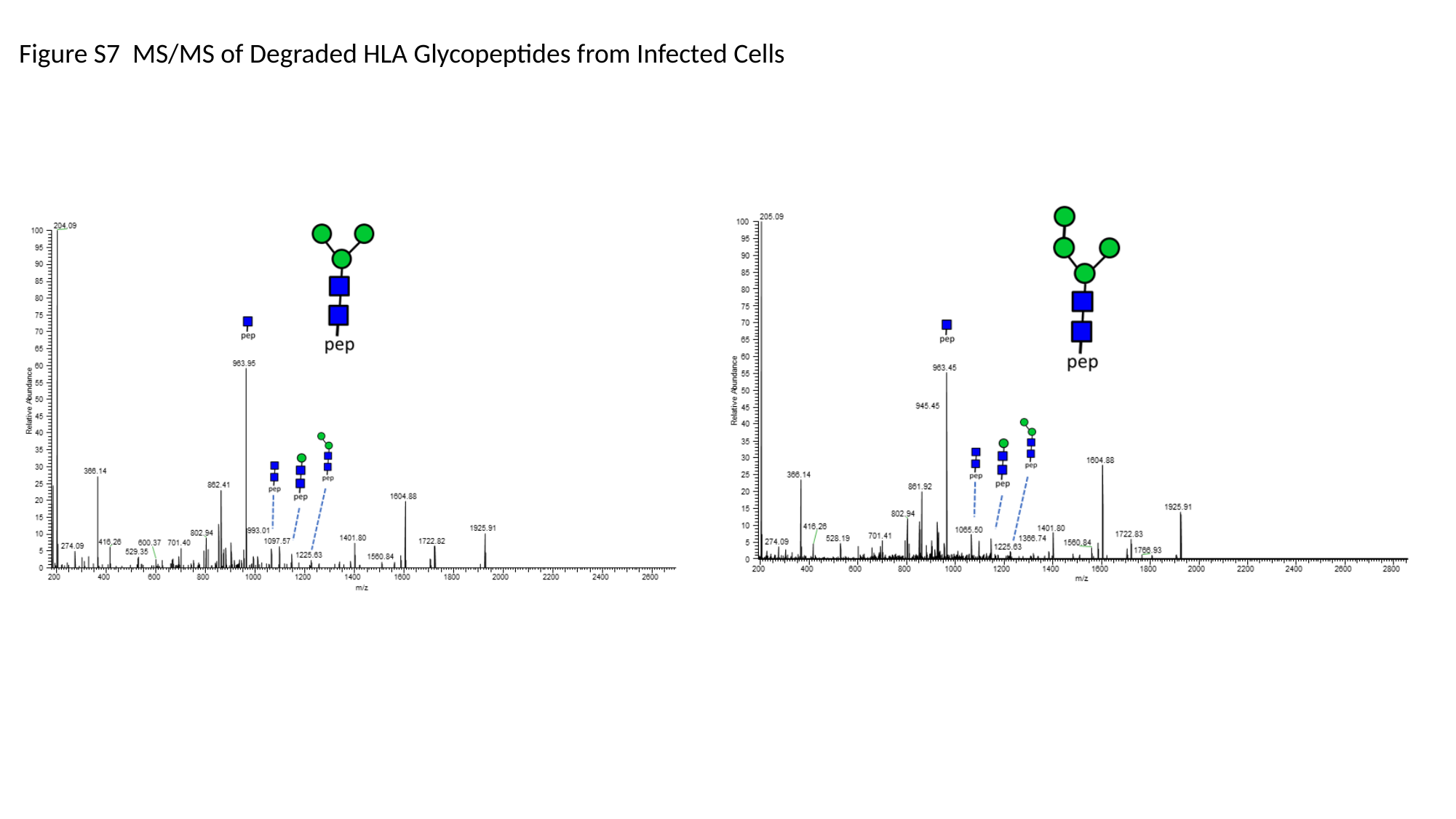

Figure S7 MS/MS of Degraded HLA Glycopeptides from Infected Cells

### Slide 8
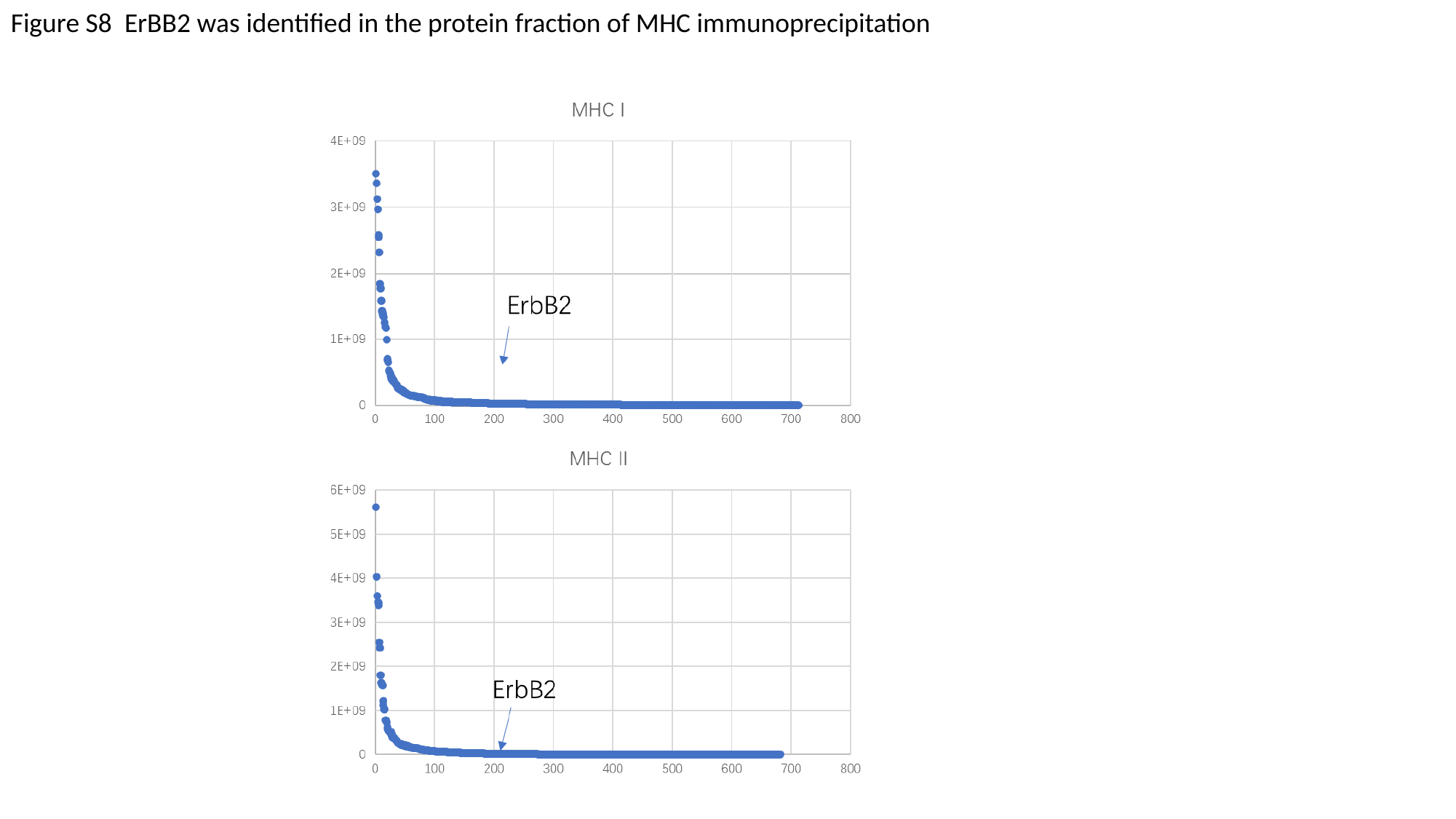

Figure S8 ErBB2 was identified in the protein fraction of MHC immunoprecipitation
